# Measurement of expression from a limited number of genes is sufficient to predict flowering time in maize

**DOI:** 10.1101/2022.12.12.520168

**Authors:** J. Vladimir Torres-Rodríguez, Guangchao Sun, Ravi V. Mural, James c. Schnable

## Abstract

Changing patterns of weather and climate are limiting breeders’ ability to conduct trials in the same environments in which their released varieties will be grown 7-10 years later. Flowering time plays a crucial role in determining regional adaptation, and mismatch between flowering time and environment can substantially impair yield. Different approaches based on genetic markers or gene expression can be used to predict flowering time before conducting large scale field evaluation and phenotyping. The more accurate prediction of a trait using genetic markers could be hindered due to all the intermediate steps (i.e. transcription, translation, epigenetic modification, and epistasis among others) connecting the trait and their genetic basics. The use of some intermediate steps as predictors could improve the accuracy of the model. Here, we are using two public gene expression (RNA-Seq) data-sets from 14-day-old-maize-seedling roots and whole-seedling tissue at v1 stage (10 day after planting) for which flowering data (days to anthesis and days to silking expressed in growing degree days) and genetic markers were also available to test the predictability of flowering time. In total, 20 different combinations between phenotypic and gene expression data-sets were evaluated. To explore prediction accuracy a random forest model was trained with the expression values of 44,303 gene models hosted in the current B73 maize reference version 5 and then the feature importance was scored based on the decrease in root mean squared error. Later several random forest models with different subsets of the most important features (genes) were trained, and this process was repeated ten times. Results from these analyses show a curve in the prediction accuracy, with an increase in the prediction accuracy as the top most important genes were added. The maximum accuracy was attained when 500 genes for whole-seedling and 100 genes for root gene expression data were used in the analysis, and thereafter adding more genes lead to a decrease in the prediction accuracy. The highest prediction accuracy using the top-most important genes was higher than that of using randomly selected whole-genome 400,000 SNPs. Finally, we described the genes controlling flowering time by looking at the most important genes in the Random forest model with the expression data from all genes. We further found MADS-transcription factor 69 (*Mads69*) using whole-seedling gene expression and the MADS-transcription factor 67 (*Mads67*) using root gene expression data, both genes previously described with effect on flowering time. Here, we aim to demonstrate the potential of selecting and using the expression of most informative genes to predict a complex trait, also to demonstrate the robustness and limitations of this analysis by using phenotypic data-sets from different environments.

## Introduction

Many traits of agronomic importance are complex in nature and are controlled by many genes with small effects. Predicting and understanding the biological control of such traits is of pivotal importance. From an agronomic or breeding perspective, predicting genotypes that will perform better in an unseen environment can help in reducing the time and efforts required in developing and releasing varieties that can perform better in new environments. The prediction of complex traits has been classically achieved by using set of genomic markers Meuwissen *et al*. (2001) termed as “genomic prediction” and a lot of progress has been made in this area Crossa *et al*. (2010, 2014); Dias *et al*. (2018); Fernandes *et al*. (2018). In the process of developing genomic prediction models a population is phenotyped and genotyped, then this population is divided in a training and testing sub-populations to test the performance of the model under a cross-validation design. One limitation of this methodology is that it does not account for various complex biological processes that exist between the genomic information (DNA sequence) and the predicted trait. Processes such as transcription, translation, post-transcriptional and post-translational regulation, coupled with some epigenetics modificatons can lead to the complexity of a trait. Models including information about some of these processes has been reported with inconclusive results.

Several studies addressing the ability of trait prediction using information related to these intermediate process has reported opposite results. In 2020, Azodi et al Azodi *et al*. (2020) compared the prediction of flowering time, yield and height of maize plants, using genomic markers and genome-wide gene expression in an association panel. They found that in general, trait prediction using genomic markers performs better than gene expression. Furthermore, selection of the most important predictors do not increase the prediction accuracy. On the other hand, Zhang *et al*., 2020 Zhang *et al*. (2020), found that using the expression of a few genes previously described with effect on maize yield can lead to better prediction than a large number of genome-wide DNA markers in a bi-parental population Zhang *et al*. (2020). A similar study was reported in cotton using contributing genes to predict fiber length and it was found that a small number of genes can predict fiber length with good accuracy in a bi-parental population Liu *et al*. (2020). Interestingly, not only the use of gene expression outperformed SNP markers in predicting several traits in mice, but also the gene expression from different tissues were found to have differential prediction accuracy Takagi *et al*. (2014) highlighting the capacity of tissue-specific gene expression to predict complex traits.

Flowering time is a complex trait controlled by the action of many genes with small effect Salvi *et al*. (2007); Buckler *et al*. (2009);Bertin *et al*. (2013); Romero Navarro *et al*. (2017), and affects several traits of agronomic importance, such as biomass and seed yield, stress avoidance and is helpful in establishing crossing strategies Jung and Müller (2009); Bendix *et al*. (2015). Flowering time also reflects the adaptation to local climate conditions Buckler *et al*. (2009) which is expected to change substantially due to global warming.

In the present study, we evaluate the prediction accuracy of two different gene expression data-sets one from root and another from the whole seedling using different flowering time as days to anthesis and days to silking, reported as growing degree days, in different phenotypic data-sets from three different environments. We used Random Forest due to its ability to capture non-linear associations between the predictors and the response variable, which is a common behaviour of “omics” data Boulesteix *et al*. (2012). Random Forest can be used to obtain two main results: the first one is a prediction model and the second to rank the features utilized to build the model. We first obtain the most-important genes to predict flowering time, then we use the expression of those genes to explore whether they increase the accuracy, lastly, we looked into those genes and found some genes previously reported as flowering time modulators in maize but also some novel genes that could potentially be involved in modulating flowering time in maize. Finally, we compared the accuracy in the prediction using gene expression with Random Forest model versus the accuracy in the prediction of genomic markers.

## Materials and methods

### Data preparation

A subset of the Wisconsin Diversity Panel (WiDiv) Mazaheri *et al*. (2019) was grown and evaluated in the field experiments conducted in Lincoln, Nebraska, as well as in East lancing, Michigan in the summer of 2020 and 2021. The flowering time data-sets from both locations in 2020 are described in Mural *et al*., 2022 Mural *et al*. (2022) while the flowering time data collected in 2021 in both locations are presented in this study (Supplemental_data_1). In brief, the experiment layout was a randomized complete block design with 2 blocks of 840 plots including repeated checks in Lincoln, NE 2020 and 2021, and 764 plots with repeated checks in East Lancing, MI 2020 and 2021. In each environment, the days to anthesis and days to silking were recorded at the plot level as the number of days from planting to the first day when 50% of plants in the plot had visible pollen shed or visible silk respectively. Each flowering time trait from each environment was further converted to respective growing degree days (GDD) as cumulative GDD from planting to flowering for each genotype. Hereafter the days to anthesis growing degree days are called anthesis and days to silking growing degree days are called silking. The trait data-set was augmented by adding additional flowering time data-sets of Peiffer *et al*. Peiffer *et al*. (2014) assembled in Mural *et al*. (2022).

### RNA-seq data-sets and gene quantification

Two sets of RNA-Seq data were used in this study. The V1 stage (approximately 10 days after planting) whole-seedling RNA-Seq data from 502 genotypes (PRJNA189400) Hirsch *et al*. (2014), and 14-day-old root RNA-Seq data on 350 genotypes (PRJNA793045), Sun *et al*. (2022). The RNA-Seq data were downloaded from NCBI using the *prefetch* and *fasterq-dump* functions inside modules SRAtoolkit v2.11 and entrez-directand v13.3. These RNA-Seq data-sets consist of paired-end samples of 100- and 75 nt lengths, respectively.

Raw reads were filtered and quality trimmed using trimmomatic v0.33 Bolger *et al*. (2014) with the following parameters: “ILLUMINACLIP:TruSeq2-PE.fa:2:30:10 LEADING:3 TRAILING:3 SLIDINGWINDOW:4:15 MINLEN:35”. Gene quantification reported as transcripts per million (TPM) was calculated using kallisto v0.46 Bray *et al*. (2016). Reads were pseudo-aligned against the version 5 Hufford *et al*. (2021) of the reference line B73 Schnable *et al*. (2009), hereafter, B73 V5 at gene level. Root gene expression followed the same procedure but libraries of some of the genotypes were duplicated, in such cases the mean was calculated for those libraries for the downstream analysis.

### SNP data-set

The Single Nucleotide Polymorphism (SNP) data-set consisting of nearly 900,000 imputed SNPs for a total of 942 genotypes involving the genotypes used in this analysis was retrieved from Mazaheri *et al*. (Mazaheri *et al*. 2019). The SNP marker data on the subset of the genotypes used in this study was further used to remove the population structure and to run the rr-BLUP model described later. To create the data frames as input in the models, common names among different data-sets were matched using the function *amatch* inside the R library stringdist Van der Loo *et al*. (2014).

### Random Forest modeling

Each of the ten phenotypic data-sets used in our analysis was screened individually. Any genotype with a phenotypic value outside 1.5 * of IQR (Interquartile range) was removed and the remaining genotypes were used in the downstream analysis. The SNP markers corresponding to each genotype in a given phenotypic data-sets were used for the Principal component analysis (PCA) using rTASSEL v0.9.28 Monier *et al*. (2022). First five principal components were used to fit a linear model as follows: *lm*(*y1 PC1* + *PC2* + *PC3* + *PC4* + *PC5*) where *y* means either anthesis, silking or gene expression. The residuals from the linear model were used as an input in the Random Forest model Breiman (2001). At first the random forest model was trained using all gene expressions as predictors and flowering time as a dependent variable using caret package Kuhn (2008) without cross-validation. Predictors (genes) were ranked based on their calculated importance, which was estimated as the decrease in root mean squared error. Then, several Random Forest models were trained using different subsets of the ranked genes (example: 10,25,50 and so on in the most important genes) and the prediction accuracy on a ten-fold cross-validation was saved. For all these models described, the number of trees was 500, and the node size was equal to 5. The number of predictors to use at each split were calculated automatically by the caret package based on the mean squared error. Correlation values were calculated with the function cor.test and then squared. All the process above was independently repeated ten times. Analysis were performed in r Team *et al*. (2013) using R studio interface.

### Genes of importance to predict flowering time

The top 500 genes in the case of whole-seedling expression data-sets and top 100 in the case of root expression data-sets were selected from the first Random Forest model on each phenotypic data-set. The Random Forest model was run for ten times and if a gene was present among all ten different independent runs the gene was considered as a candidate to control that trait. Description to the genes was added based on the Interproscan description found in the Maize Genomic Data Base, MaizeGDB (https://download.maizegdb.org/Zm-B73-REFERENCE-NAM-5.0/, Woodhouse *et al*. (2021)) and an updated classical maize genes list Schnable and Freeling (2011)

### rr-BLUP modeling

A subset of 400K SNPs were randomly selected from the original SNPs data-set (see Data preparation) using the function −*subsetSites* inside tassel TASSEL v5.2 Bradbury *et al*. (2007). The data frame with the phenotypic information and the SNPs for the corresponding genotypes was created for each location. The function *mixed.solve* inside the rrBLUP library Endelman (2011) was used to estimate the marker’s effects. The first five Principal component values from a PCA analysis in rTASSEL v0.9.28 Monier *et al*. (2022) using 400k randomly selected SNPs and only the genotypes in the analysis were included as a fixed effect in the model. The accuracy of the model was tested in five-fold cross-validation.

### Data and code availability

Raw flowering time and raw results generated as a part of this study are provided as supplemental data. Any additional data and codes used in the analysis will be made available on request.

## Results

### Flowering time and Gene expression data-sets

In order to explore the ability of gene expression data-sets to predict flowering time (anthesis and silking), we used two different gene expression data-sets; one from whole-maize seedling at V1 stage Hirsch *et al*. (2014) and another from roots of maize plants at 14 days after planting Sun *et al*. (2022). These gene expression data-sets include different subsets of the expanded 942-WiDiv panel Mazaheri *et al*. (2019). These data-sets were matched with flowering time data in growing degrees days, collected across different locations many of these data are summarized in Mural *et al*. Mural *et al*. (2022), but the ones collected from NE2021 and MI2021 are reported here for the first time. A total of 5 different environments were used as part of this study, which include data from Nebraska and Michigan collected in 2020 and 2021. Flowering time data reported as Peiffer2014 was the mean of values collected from New York, Missouri, and North Caroline Mural *et al*. (2022); Peiffer *et al*. (2014). In general we have more genotypes matching the whole-seedling gene expression data-set than genotypes matching the root gene expression with an average of 400 and 175 genotypes, respectively. The phenotypic data-set with more number of genotypes matching with both expression data-sets was present in Peiffer2014 (438 in whole-seedling expression data-set and 181 in root expression data-set) while the one with the least number of genotypes was NE2020 with 393 and 176 genotypes matching seedling and root expression data-sets respectively. The number of genotypes used in each analysis are presented in Supplemental table S1. The distribution of anthesis and silking varies in each phenotypic data-set as well as the maximum and minimum values. The distribution of different flowering time data-sets after outlier removal are shown in figure 1.

**Figure 1.**
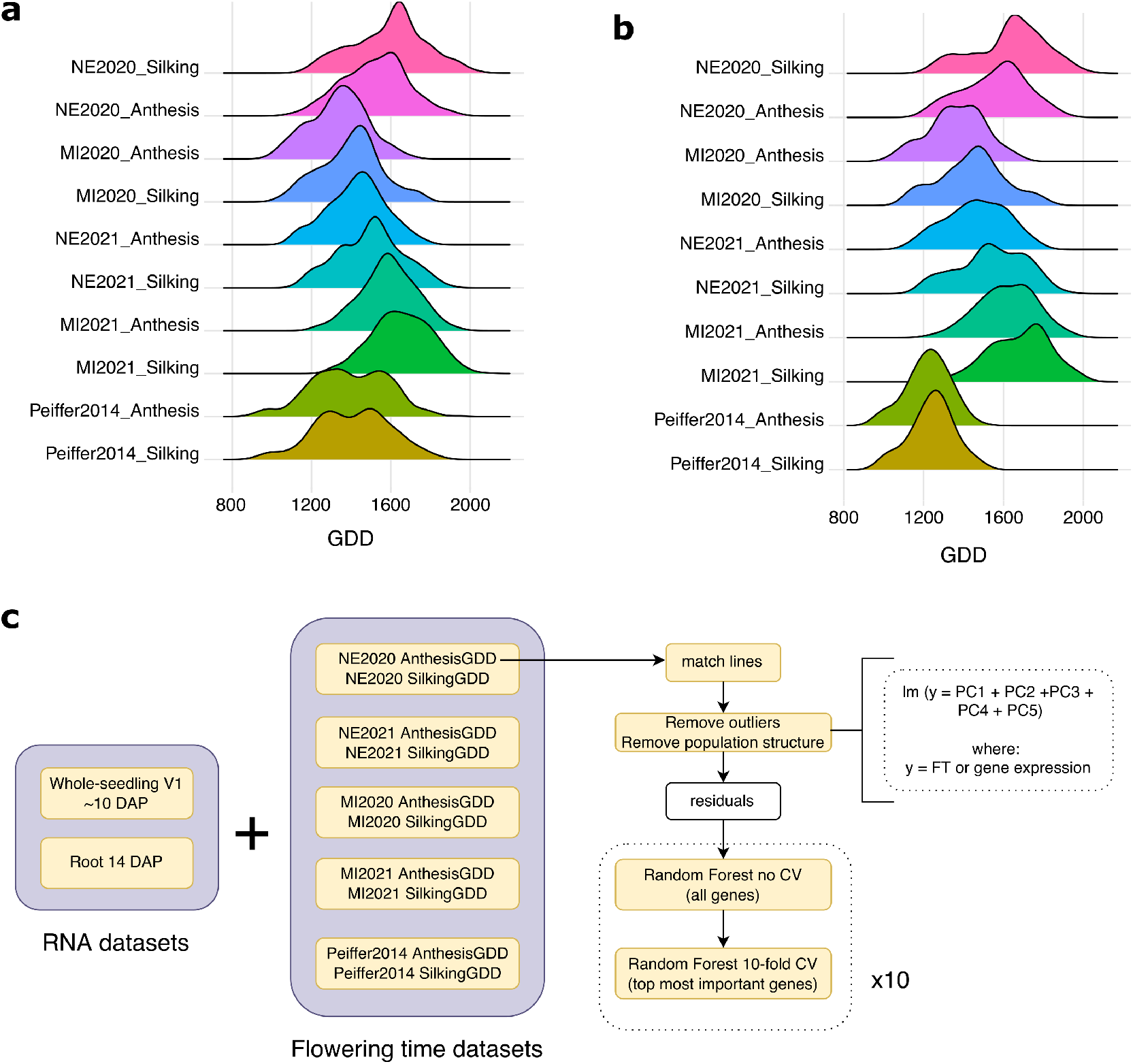
Data-sets used in this study and the general workflow. Distribution of days to anthesis and days to silking in growing degree days among the data-sets used with the whole-seedling (**a**) and with the root (**b**) gene expression data. **c** General Diagram of the process used to predict accuracy from different phenotypic data-sets. DAP stands for days after planting and GDD for growing degree days

### Using the expression of most informative genes can improve prediction accuracy

Each trait data-set was pre-processed by removing outliers to ensure that the downstream analysis is not biased/affected by any obvious outliers. Apart from extreme outliers, population structure can also affect the ability of the model performance and affects the prediction ability across populations. To address this we ran a principal component analysis on both the dependent variable (each phenotypic trait) and independent variables (gene expression) using the SNP markers corresponding to each genotype in a given phenotype. The first five principal components were extracted for each data-set and were further used to fit a linear model. The resulting residuals from the linear model were further used as input in the random forest (RF) model.

Running the random forest model ten times with 10-fold cross-validation using gene expression of all gene models published in the B73 V5 reference genome led to a variable prediction accuracy of the phenotypic data-sets in different environments. In general, the mean phenotypic prediction accuracy was slightly higher when expression data from the whole seedling was used and was slightly lower in some cases when root expression data was used (figure 2a and b). For both anthesis and silking the highest prediction accuracy was observed for NE2020 (average *R*^2^ of then reps = 0.41 [seedling] and 0.36 [root]) while the lowest prediction accuracy was observed for MI2021 using whole-seedling and root gene expression data in the analysis (average *R*^2^ of then reps = 0.27 and 0.12, respectively) (figure 2a and b). We also observed consistency between the prediction accuracy of anthesis and silking in each environment in both cases where either whole-seedling or root expression data was used in the prediction model.

**Figure 2.**
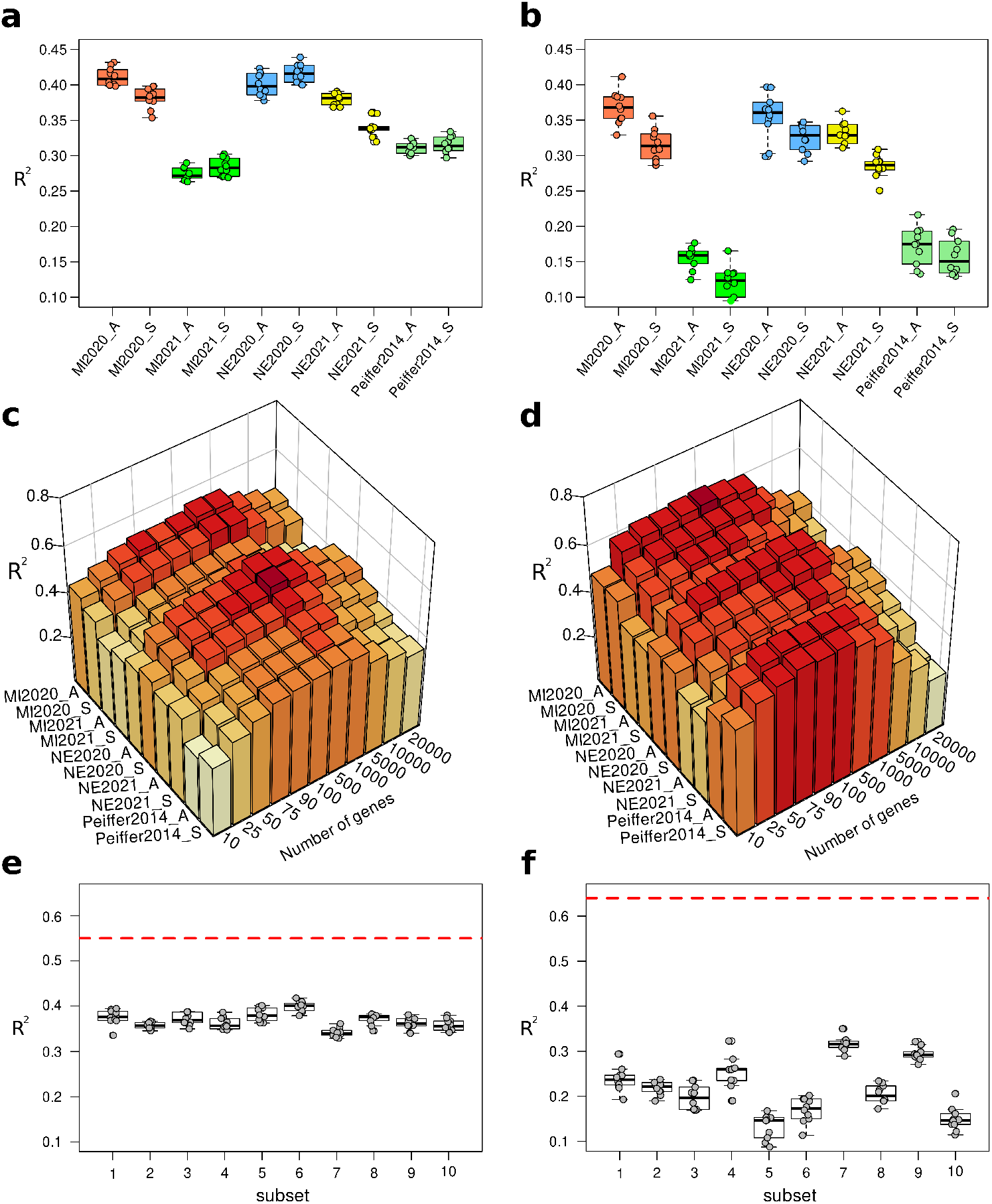
Flowering time prediction accuracy across different data-sets. Prediction accuracy of days to anthesis and days to silking expressed in growing degree days among different data-sets using the whole-seedling **(a)** and with the root **(b)** gene expression data. **c** 3D histograms representing the *R*^2^ values for different flowering time subsets using the expression of different number of top-most-important genes in whole-seedling and **d** root gene expression data-sets. e Prediction accuracy of anthesis in NE2020 using the expression of 500 randomly selected whole-seedling genes. The red dotted line represents the prediction accuracy of NE2020 using the expression of the 500 top-most-important root genes. Subsets in the x-axis means 10 different subsets of genes from the original file. f Prediction accuracy of anthesis in NE2020 using the expression of 100 randomly selected root genes. The red dotted line represents the prediction accuracy of NE2020 using the expression of the 100 top-most-important root genes. Subsets in the x-axis means 10 different subsets of genes from the original file. Boxplots in a,b,e and f, represent data from 10 independent Random Forest modeling in a 10-fold cross-validation. 3D histograms are showing the mean value of 10 independent Random Forest modeling in a 10-fold cross-validation. In sub-panels a-d, in the phenotypic data-set, “A” means anthesis and “S” means silking.

Fitting models with the gene expression of informative genes instead of gene expression from all the genes are reported to improve the prediction accuracy Zhang *et al*. (2020); Liu *et al*. (2020). Thus, we further selected the gene expression from most informative genes in our prediction model. To detect the most important features or genes in this case we ran a RF model without cross-validation and ranked the genes based on their importance, which is defined as the decrease in the root mean squared error. We then selected different subsets (10, 25, 50, 75, 90, 100, 500, 1000, 5000, 10000, and 20000) of these most-important genes and tested the prediction accuracy of anthesis and silking. This led to an increase in the prediction accuracy in all phenotypic data-sets included in both the cases when whole-seedling and root gene expression data-sets were used in our prediction model. Maximum prediction accuracy was attained when the expression data from 500 top-most-important genes were used in the case of seedling and 100 top-most-important genes in the case of root gene expression data-set (figure 2c and d). Further addition of genes tend to stabilize and did not increase the prediction accuracy. In fact we observed that the accuracy decrease after a certain point. We observed decrease in the prediction accuracy when more than 1000 genes were used.

To rule out the possibility that the maximum accuracy is due to the number of genes and not their expression, we randomly selected 500 genes (whole-seedling expression data) ten times and ran 10-fold cross-validation RF model for ten different times (for the same subset of 500 genes) using NE2020_Anthesis data as a dependent variable. The prediction accuracy was consistently lower in all 10 cases (figure 2e) compared to the prediction accuracy obtained with the expression of the top-most-important genes (average *R*^2^ of then reps = 0.55) (figure 2c). We repeated same procedure using 100 randomly selected genes from the root data-set and NE2020_Anthesis as a dependent variable. Similar to whole-seedling expression data, the prediction of these data-sets was also lower (figure 2f) when compared to the one with the top-most-important genes (average *R*^2^ of then reps = 0.63)(figure 2d). Together these results suggest that it is possible to increase the prediction accuracy of flowering time using the expression of a few selected genes.

Because the original whole-seedling gene expression data-set was composed of 502 genotypes and the root gene expression data-set was composed of 350 genotypes, the number of genotypes used to calculate the prediction accuracy was larger in the case of whole-seedling expression data than the root gene expression data S1. This opens up a question of whether the differences in the accuracy are due to the number of genotypes used in our model or because of the expression of the genes. To explore this we ran an RF model using the exact same genotypes for both gene expression data-sets.

We observe that the prediction accuracy was not just affected by the origin of the tissue (whole-seedling or only root) used to get the gene expression but also the genotypes we used. The ability of root expression data to predict the phenotype with higher accuracy while using the top most important genes was still observed but was not consistent across the phenotype as observed earlier. In MI2021 (average *R*^2^ of then reps at highest value = 0.53) and NE2021 (average *R*^2^ of then reps at highest value = 0.60), prediction accuracy was higher when we used whole-seedling expression data, while, using root expression data, only Peiffer2014 had higher prediction accuracy (average *R*^2^ of then reps =) than the same genotypes with root gene expression. For the remaining environments, the prediction accuracy was the same. Interestingly, in some cases, like NE2020 and MI2020, the accuracy was higher if compared against the one with all genotypes from the phenotypic data-set (Figure 3).

**Figure 3.**
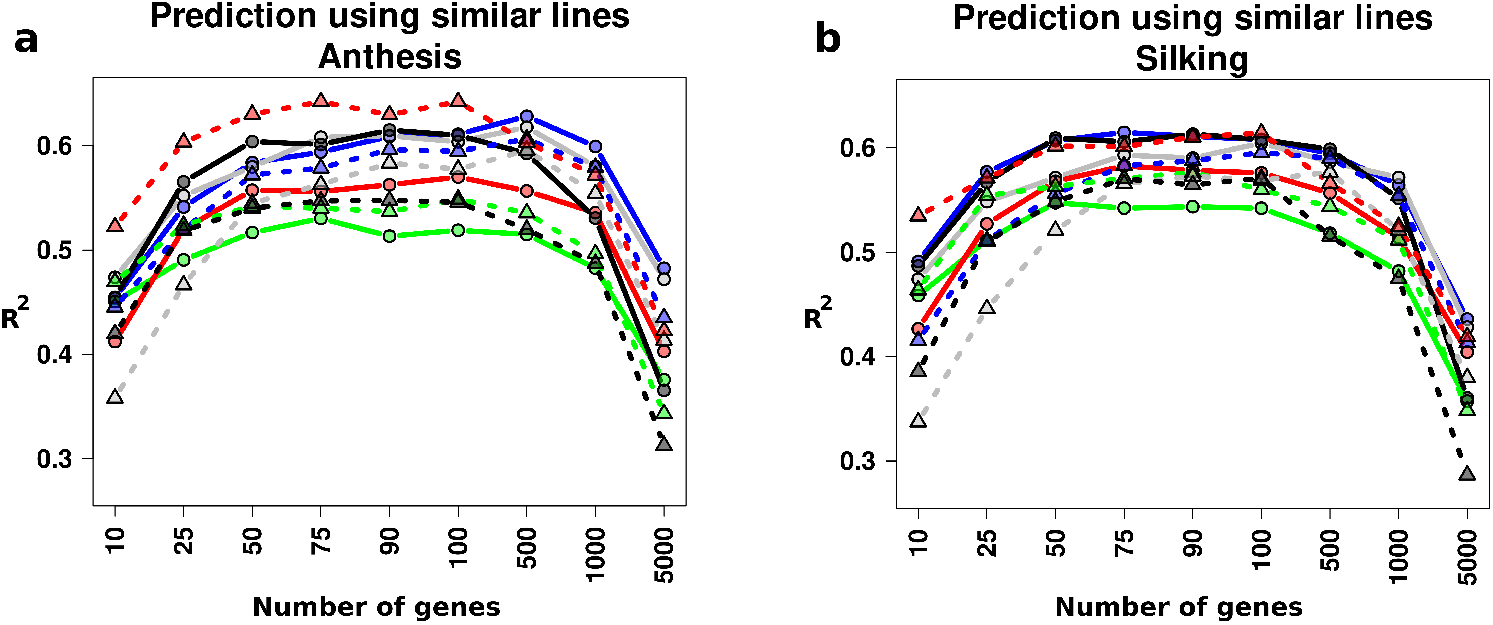
Prediction accuracy using the same genotypes from each gene expression data-set. Prediction accuracy of anthesis **(a)** and silking **(b)** using the same genotypes for each phenotypic data-set. Circles (solid line) and triangles (dotted line) represent root and whole-seedling gene expression data, respectively. Color code is as follows: Blue = MI2020, green = MI2021, grey = NE2020, red = NE2021 and black = Peiffer2014.

### Top-most important genes are found across different data-sets in whole-seedling and root gene expression data

We have shown above that the prediction accuracy can increase when we use the expression of the top-most-important genes. To explore which are the genes defining anthesis and silking in each phenotypic data-set, we extracted the top 500 genes for each data-set (for the 10 independent runs) in the whole-seedling and root gene expression data-sets. We found a non so small number of genes for each phenotypic data-set. Using the whole-seedling gene expression data-set the average of different genes per phenotypic data-set were 2882 while using the root data were 3273 (supplemental_data_2 and supplemental_data_3). To select the best candidates we detected genes that show consistency among the 10 different runs (i.e., genes present in each of the ten runs). In total there are 397 genes in whole-seedling expression and 191 genes in root expression present 10 times across all different phenotypic data-sets, with some genes found in more number of data-sets than others (supplemental_data_2 and supplemental_data_3). The most common genes for phenotypic data-sets using whole-seedling gene expression as predictors, with more than six out of ten appearances, are Zm00001eb143080(8) and Zm00001eb042250 (8) (Fig. 4a), while using root data, the genes are Zm00001eb412060 (7) and Zm00001eb424860 (6) (Fig. 4b). Interestingly, the gene model Zm00001eb143080 correspond to the MADS (MCM1/AGAMOUS/DEFICIENS/SRF) −transcription factor 69 (*Mads69*). For important genes with less appearances the Interproscan description includes “*Domain of unknown function DUF676*”, a “*lipase-like*” (Zm00001eb214270), a *UDP-glucosyltransferase* (Zm00001eb310200 [root data]), moreover, we found another MADS-transcription factor, *Mads67* (Zm00001eb327040), using root gene expression as predictors in six out of ten phenotypic data-sets. Full list of genes is provided as supplemental_data_2 and supplemental_data_3. Expression of some of these genes are shown in Figure 4c-e. Although small, a negative correlation is observe between the MADS-Box transcriptions factors and the flowering time. It is noteworthy that one of the most common gene found using whole-seedling gene expression (Zm00001eb04225) is also found using root gene data, and is the only gene found in both expression data-set. We observe differences in the number of genes that each phenotypic data-set has. Both NE2020_Anthesis and NE2020_Silking had all 12 genes which were most consistent among all phenotypic data-sets using both gene expression data-sets (Fig. 4a-b); In contrast, Peiffer2014_Anthesis and Peiffer2014_Silking does not have any gene among the 12 top most consistent genes using the whole-seedling gene expression, similarly none of the consistent genes were found for MI2021 using root gene expression data.

**Figure 4.**
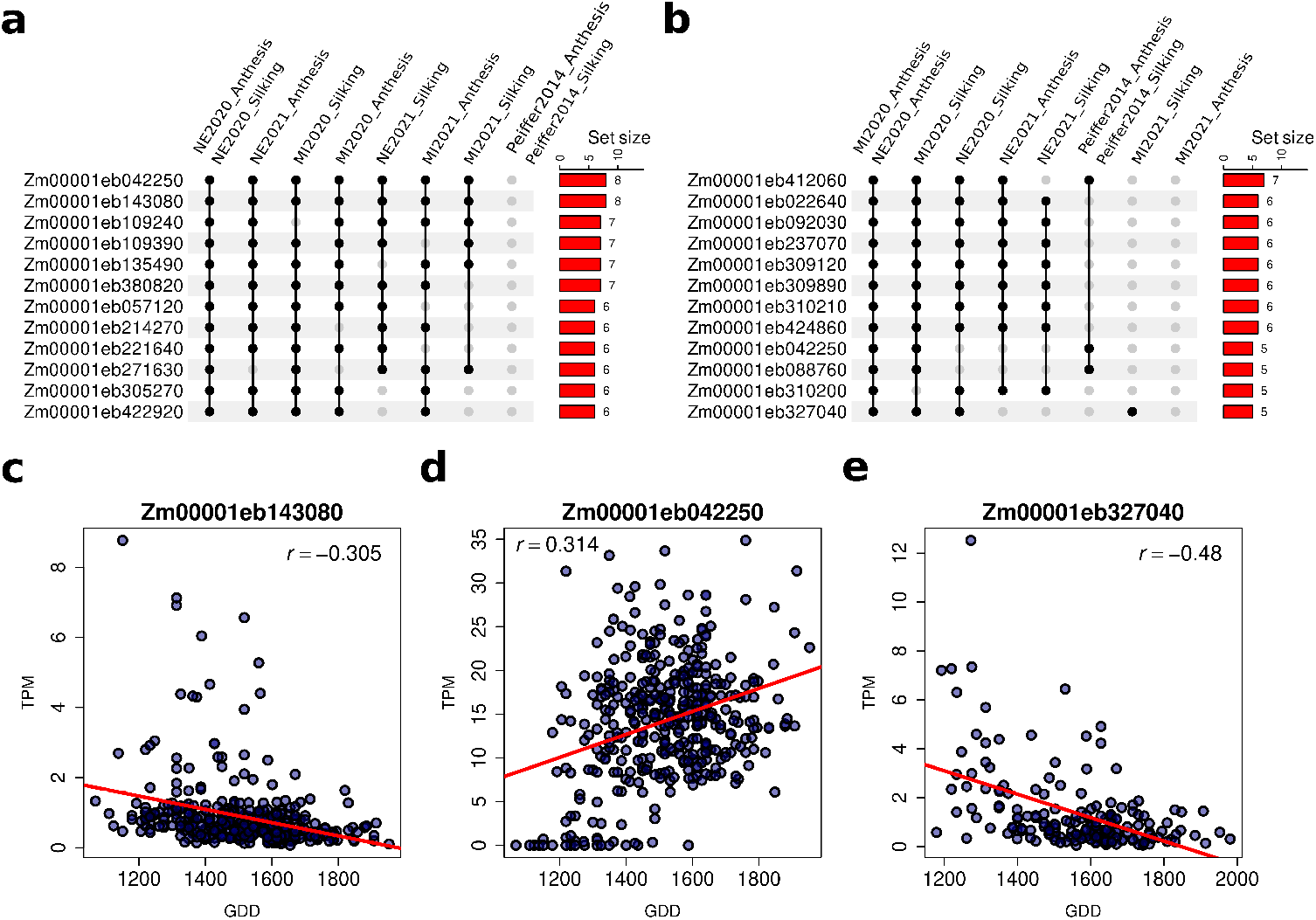
Genes found inside the top-most-important genes across different sub-sets. **a** 12 most consistent genes recovered from the 500 top-most important genes using the whole-seedling gene expression data across different field locations. **b** 12 most consistent genes recovered from the 500 top-most important genes using the root gene expression data across different phenotypic data-sets. Filled dot represents genes found all times as top most important. Barplots in the right side of the plot represents the numbers of phenotypic data-sets in which a gene was found

### The gene expression of the top-most important genes outperform the accuracy of genomic prediction

Classically trait prediction has been done with genomic markers Meuwissen *et al*. (2001). To explore how anthesis and silking prediction using the gene expression is compared against genomic markers (SNPs) we run rr-BLUP with 400,000 randomly selected markers on all phenotypic data-sets. The prediction accuracy with SNPs was different among phenotypic data-sets. Using phenotypic data-set from the whole-seedling analysis, MI2020 was the data-set with highest accuracy (*R*^2^ = 0.39 [anthesis] and 0.44 [silking]) and Peiffer2014 was the one with the lowest accuracy (*R*^2^ = 0.02 [anthesis] and 0.03 [silking]) (Figure 5a). When we used root gene expression data, the highest accuracy is for NE2020 anthesis and silking whereas the lowest is for Peiffer2014 (Figure 5b). Although small differences, prediction using SNPs (with rr-BLUP) is similar to use all genes (with RF). In some cases prediction is higher with SNPs for example MI2021, but in others like Peiffer2014 or NE2021 the prediction is better when we use gene expression and Random Forest. Importantly, the prediction is always higher when we use the top most important genes, either 500 or 100, for whole seedling and root data-sets, respectively.

**Figure 5.**
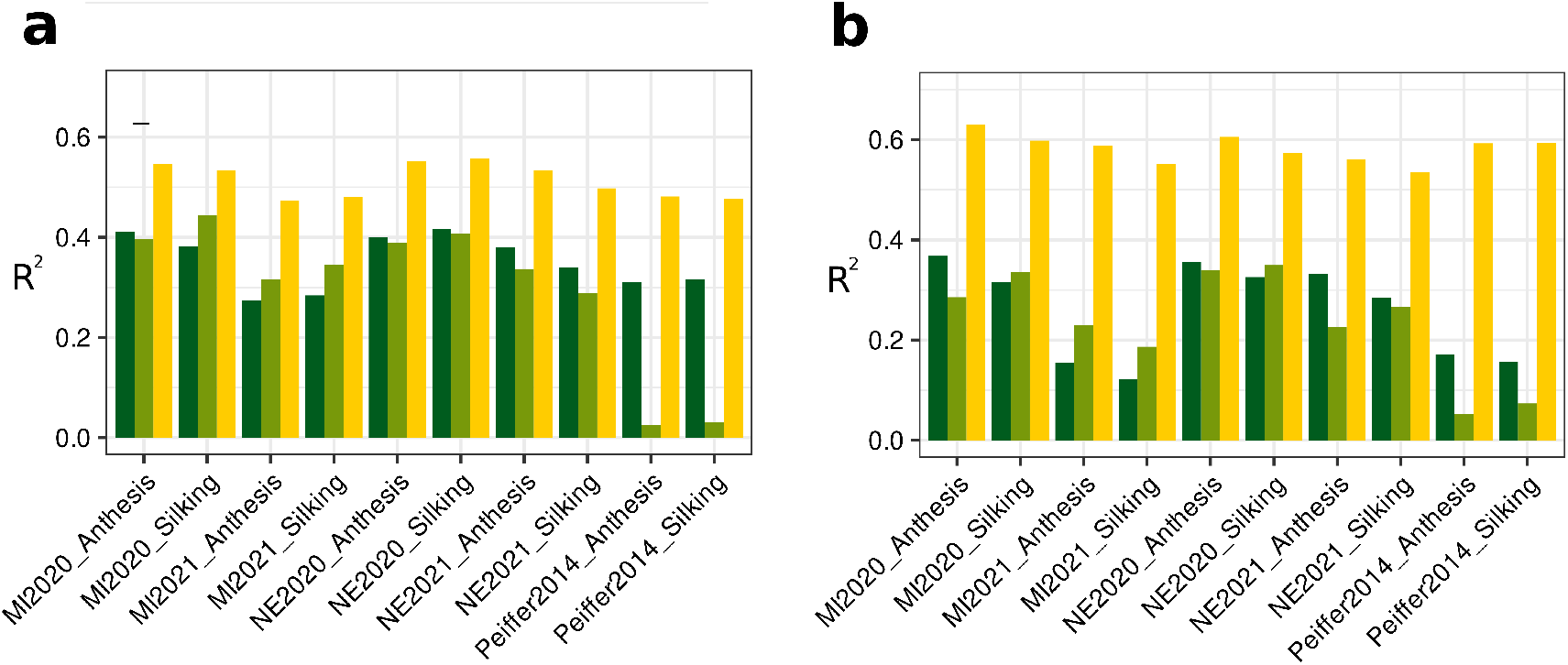
Comparison of prediction accuracy using SNPs and gene expression. **a** Barplots showing the *R*^2^ value for the different data-sets used in the whole-seedling analysis. Dark green = expression of 44,303 genes, light green = 400,000 random SNPs across the genome, and yellow = expression of 500 top most-important genes. **b** Barplots showing the *R*^2^ value for the different data-sets used in the root analysis.Dark green = expression of 44,303 genes, light green = 400,000 random SNPs across the genome, and yellow = expression of 500 top most-important genes.

## Discussion

Trait prediction can help breeders and researchers save time, money and resources by estimating the possible value of a given trait without the necessity of growing and phenotyping all genotypes under consideration. Breeders often deal with complex traits which are controlled by many genes. Flowering time is a complex trait with a direct effect on yield that is controlled by many genes Romero Navarro *et al*. (2017). Here we evaluate the prediction accuracy of flowering time (Days to anthesis and days to silking expressed as growing degree days) in maize from gene expression using Random Forest. In order to do that, we use different sub-sets of the WiDiv panel, with published Mural *et al*. (2022) and unpublished flowering time as phenotypic data-sets and two publicly available gene expression data-sets derived from RNA-Seq Hirsch *et al*. (2014); Sun *et al*. (2022).

One of the drawbacks while working with a population and using cross-validation scheme to test the model performance is that the population structure can affect the predicted value Werner *et al*. (2020) If the population structure is not considered and is not corrected/ not sufficiently corrected then the risk of finding false positives increases substantially Larsson *et al*. (2013). Since our goal was not only to explore the ability to predict flowering time using gene expression data but also to pinpoint the genes that best describe it, we removed some population structure from our analysis by fitting a linear model with the first five principal components calculated from a PCA with the SNPs corresponding to the genotypes analyzed in each data-set.

One plausible advantage of our study is the use of several phenotypic data-sets each including not only different number of genotypes but also different number of overlapping genotypes. Using whole-seedling gene expression, we observed that the accuracy can vary across phenotypic data-set 2, with accuracy almost 150% higher than the lowest one, this variation is higher (230%) when we used root gene expression. This points out the importance of using different data-sets to give a better sense of the ability of the technique used. When using all the genes, the higher accuracy reported as *R*^2^ was around 0.42 for NE2020_anthesis while the lowest was around 0.28 for MI2021, this values are similar to the one reported in Azodi *et al*., 2020 for flowering time Azodi *et al*. (2020) with a value of 0.25 (calculated from a Pearson correlation 0.5) using Random Forest and the same original whole-seedling trasncriptomic data.

Previous studies have suggested that using the expression of the contributing genes can increase the prediction accuracy of complex traits Liu *et al*. (2020); Zhang *et al*. (2020). We have defined which are the genes modulating days to anthesis and days to silking by ranking genes based on their importance as depicted based of the decrease in the root mean squared error. To define which are the most important genes we fitted several Random Forest models using different sub-sets of genes, interestingly the number of genes required to achieve the maximum prediction accuracy across phenotypic data-sets was similar for each gene expression data-set (500 genes in case of whole-seedling and 100 in case of root expression data). We ruled out the possibility that this is because of the number of genes rather than the expression of the genes themselves by running the same pipeline with 500 (whole-seedling expression data) and 100 (root expression data) randomly selected genes as a negative control.

One advantage of working with gene expression rather than genomic markers is that we are working at a gene level, which facilitates the identification of candidate genes modulating the trait under investigation. An average of 2800 and 3200 genes were classified as important genes when we used whole-seedling expression and when root expression, reflecting the complexity of the trait, in an attempt to select which are the best genes describing flowering time we selected genes that were present in every single run (10 times). We found genes matching the condition above in several phenotypic data-set, one of the most common gene using whole-seedling expression data was Zm00001eb143080, corresponding to *Mads69*. Over-expression of *ZmMADS69* causes early flowering, while a transposon insertion mutant of *ZmMADS69* exhibits delayed flowering Liang *et al*. (2019). Consistently with this result, we observed that genotypes with higher growing degree days have lower levels of *ZmMADS69* (Figure 4c). Another gene reported with a direct effect on flowering time is *Mads67*, Zm00001eb327040, knocking out *ZmMADS67* using CRISPR/Cas9 delays flowering time in maize Sun *et al*. (2020), indicating that they promote floral transition. These results are in agreement with the expression levels of *ZmMADS67* in our gene expression data-set. Genotype that needed more growing degree days to flower have less expression (Figure 4e). It is noteworthy that here we are reporting the finding of *Mads67* in root gene expression data of maize plants at 14 days after planting which provide new insights of the complexity of flowering time and suggest the involvement of root system at early stages to define flowering time in later stages of the plant. A similar result was reported for *Rap2.7* which is not only controlling flowering time but also the number of the whole-seedling-borne brace roots in maize Salvi *et al*. (2007);Buckler *et al*. (2009);Li *et al*. (2019).

Several MADs-box transcription factors has been described with roles in root development, for example *OsMADS25* promotes the growth of primary root and lateral roots Zhang *et al*. (2018), and *OsMADS57* promotes root elongation Huang *et al*. (2019). Interestingly, both genes show function in a nitrogen-dependent manner. No information was found depicting direct role of *Mads67* in root development. In our study we notice that the expression levels of *Mads67* in plants at 14 days after planting is influencing flowering time. However, more studies are needed to explore this mechanism. Only one gene was found ten times as an important gene using both gene expression data-sets, Zm00001eb042250, a gene coding for UDP-glucosyltransferase. Although no direct information linking flowering time and the actions of UDP-glucosyltransferases was found, two different hypothesis can explain this gene among others UDP-glucosyltransferases as the top most important genes in different sub-sets. UDP-glucosyltransferases are a superfamily of enzymes that transfer a glycosyl group from a UDP-sugar to a small hydrophobic molecule, a process called glycosylation. The principal donor of the glycosyl group is UDP-glucose but also UDP-galactose, UDP-xylose and UDP-ramnose can act in this process Cho *et al*. (2018). Thus, 1) UDP-glucosyltransferases are affecting the levels of sugar inside some organs and its been stated that flowering time is affected by sugar balances Cho *et al*. (2018) and 2) glycosylation controls levels of endogenous gibberellins, which has been reported to control flowering time Yamaguchi (2008); Ostrowski and Jakubowska (2014).

Initially, we used different subset of genotypes to predict flowering time, for most phenotypic data-sets we have twice the number of genotypes we used with the whole-seedling gene data compared to the root gene expression data. To explore if this difference has an impact in prediction accuracy, we fit Random Forest models using the same genotypes and only changing the origin of gene expression (whole-seedling or root tissue). We observe some differences compared to our initial results with all possible genotypes. First, the trend of root gene expression predicting better than whole-seedling data was not obvious. For some phenotypic data-sets prediction accuracy was better when we used root data but this could be explained by other factors like the genotypes used or the environment in which the original phenotypic data-set was collected. Again, these results point at the importance of evaluating the proposed technique using different data-sets from different environments, which in some case could be difficult due to the cost of growing genotypes in different fields or different years. One key difference we noted was the capability of the model to find candidate genes, whereas in the first case using all possible genotypes the number of genes present in ten times independent runs across different data-sets was 397 and 191, for whole-seedling and root data, respectively. Lower number of genes were found when less number of genotypes (143 & 98) were used, this could be explained by the decrease in the information we are providing in our model as input leading to a lack of statistical power. Therefore based on these results, if the end goal is to find candidate genes modulating a given trait we suggest to use a good number of genotypes for the analysis, on the other hand if the end goal is just to predict a trait fewer genotypes could be sufficient.

## Conclusion

It is possible to estimate the flowering time (anthesis and silking), using genome-wide gene expression data and the prediction accuracy can be substantially increased by selecting most important genes detected by our random forest model. The higher prediction accuracy using the most important genes based on expression data was even higher then the prediction accuracy obtained when the SNP markers were used for the prediction. Both the expression data used in this study were from two different organs at the seedling stage and from the different field conditions in which the phenotypic traits were tested. It is important to note the possibility of predicting flowering time with higher accuracy from the expression data obtained from different tissue collected from different fields with different conditions. This opens up the possibility of selecting tissue from a different organ or from a particular developmental stage in order to increase prediction accuracy. Here we worked with Flowering time because of its relation with yield and availability of phenotypic data on large set of lines grown in different environments. In the future it will be interesting to explore how different tissues or development stages are capable of predicting not just yield but also other economical important traits like plant height or root development.

## Acknowledgments

This work was completed utilizing the Holland Computing Center of the University of Nebraska, which receives support from the Nebraska Research Initiative. Special thanks to members of the Schnable lab who kindly collect flowering time data.

## Author contributions

J.C.S and J.V.T.R conceived the study. G.S. conducted experiments and generated data. J.V.T.R conducted analyses and visualized the results. J.V.T.R and R.V.M authored the initial draft of the manuscript. All authors contributed to writing and editing and approved the final version of the manuscript.

---

**Table S1.** Number of genotypes used from different fields

| Data-set | Whole-seedling RNA | root RNA | genotypes in both RNA |
| --- | --- | --- | --- |
| NE2020_Anthesis | 393 | 176 | 128 |
| NE2020_Silking | 393 | 176 | 128 |
| MI2020_Anthesis | 399 | 178 | 130 |
| MI2020_Silking | 388 | 176 | 128 |
| NE2021_Anthesis | 412 | 185 | 136 |
| NE2021_Silking | 413 | 185 | 136 |
| MI2021_Anthesis | 400 | 171 | 126 |
| MI2021_Silking | 402 | 171 | 126 |
| Peiffer2014_Anthesis | 438 | 181 | 133 |
| Peiffer2014_Silking | 434 | 182 | 133 |

